# constclust: Consistent Clusters for scRNA-seq

**DOI:** 10.1101/2020.12.08.417105

**Authors:** Isaac Virshup, Jarny Choi, Kim-Anh Lê Cao, Christine A Wells

## Abstract

Unsupervised clustering to identify distinct cell types is a crucial step in the analysis of scRNA-seq data. Current clustering methods are dependent on a number of parameters whose effect on the resulting solution’s accuracy and reproducibility are poorly understood. The adjustment of clustering parameters is therefore ad-hoc, with most users deviating minimally from default settings. `constclust` is a novel meta-clustering method based on the idea that if the data contains distinct populations which a clustering method can identify, meaningful clusters should be robust to small changes in the parameters used to derive them. By reconciling solutions from a clustering method over multiple parameters, we can identify locally robust clusters of cells and their corresponding regions of parameter space. Rather than assigning cells to a single partition of the data set, this approach allows for discovery of discrete groups of cells which can correspond to the multiple levels of cellular identity. Additionally `constclust` requires significantly fewer computational resources than current consensus clustering methods for scRNA-seq data. We demonstrate the utility, accuracy, and performance of `constclust` as part of the analysis workflow. `constclust` is available at https://github.com/ivirshup/constclust^1^.

## 2 Introduction

Single cell RNA sequencing (scRNA-seq) provides a snapshot of one of the fundamental units of biology: the cell itself. The molecular scale of scRNA-seq promises to unlock new dimensions to the developmental or anatomical cell catalogue - that of cell type specificity within a tissue. This improved resolution of cellular states may encompass the identification of new discrete cell types, or alternatively illustrate a spectrum of phenotypes spanning cellular responses to environmental, circadian or other molecular cues. One of the things that we aim to do with this new view is enumerate and catalogue all of the of cells found within an organism.

A catalogue of all cell types and cell states assumes that cells sharing molecular attributes will also share other attributes, such as lineage or function. This is a reasonable assumption evidenced by decades of cell profiling - from antibody-based methods for classifying cell types, to transcriptome profiling of cell populations [1]. Traditionally, molecular profiling of groups of cells relies on prior identification of the classes of cell being introduced into the comparison. In contrast scRNAseq relies on unsupervised clustering methods to identify discrete cell groups. Unsupervised clustering methods aim to partition cells on their expression profiles [2]. Further analyses such as the identification of unique marker genes or expression of pathways, rely on an accurate clustering, and classification of cluster members, so it’s critical that this part of the analysis is rigorous and reproducible. Unfortunately, most clustering methods are neither.

Despite being a popular and important part of many analyses of scRNA-seq data, the best practice for partitioning of transcriptomes into cell types is far from settled. In fact, despite most popular clustering methods producing a flat clustering (e.g. where each sample is assigned to one partition of the data set), there is broad recognition that we’re failing to capture the true structure of the data. We view cells types as being both continuous and multilevel: we can classify cells as part of a developmental hierarchy or as their resulting cell state, which may not have complete correspondence with each other. A well studied example of this is the myeloid lineage, where monocytes and macrophages can arise from different ontogeny but share functional characteristics [3]. This ability to find multilevel structure in single cell datasets described as an overarching theme of a set of grand challenges in the field [4].

While methods that group cells into a single flat partitioning cannot capture multiple levels of structure within a single solution, there have been a few attempts to address this by varying the hyper-parameters used (e.g. parameters the user chooses). For example, `clustree` lets a user visualize the relationships between clustering solutions as a hyper-parameter varies [5]. Similarly in the trajectory method PAGA, clusterings from different levels of resolution are used to model the data set at different levels of granularity [6]. These methods provide ways to interrogate the multilevel structure of the data through parameter choice. However, the responsibility is still on the user to assess the quality of the representation.

One of the most popular clustering methods for scRNA-seq is modularity maximization based community detection on nearest neighbor graph of the measured transcriptomes. This finds clusters of cells withing the neighbor graph by finding groups of cells which have more edges connecting them than would be expected under a null model [7]. While a variety of implementations are used with single cell data the method is often referred to as “Seurat” clustering in the literature [8] [9]. In comparisons of methods for unsupervised clustering of scRNA-seq data this method is frequently highly ranked for both speed and accuracy [10] [11]. Despite this, results can be unstable under small changes in parameter values Fig. 1. An important parameter for this class of method is the resolution parameter (*γ*) – varied by both PAGA and `clustree`. A systematic way to choose the value of *γ* has yet to be addressed in the single cell literature, although it has been studied in the community detection literature.

**Figure 1:**
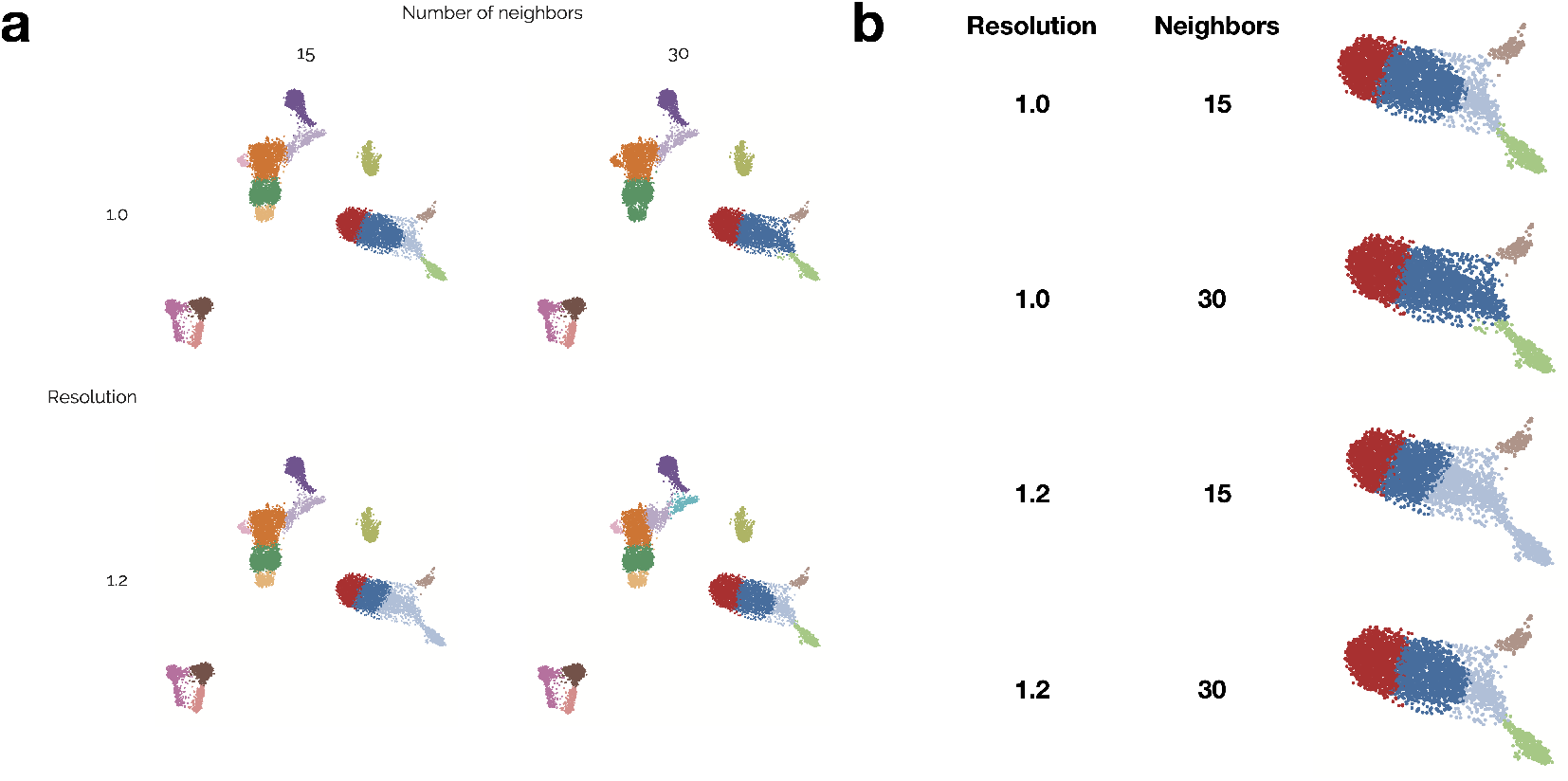
Varying clustering hyperparameters demonstrates common clustering challenges. (A) Four different clustering solutions of peripheral blood mononuclear cells (PBMCs) generated near default settings using the ‘leiden’ algorithm as used by Scanpy. Clockwise from top left, parameters are: 15 neighbors – resolution of 1, 30 neighbors – resolution of 1, 30 neighbors – resolution of 1.2, and 15 neighbors – resolution of 1.2. (B) highlights a subset of the data set with highly variable cluster membership.

Community detection is the problem of finding groups of highly connected nodes within a network. One of the most widely used approaches from this field is modularity optimization. Modularity is a measure of how interconnected each group is when compared to a null model, where connectivity is randomly distributed. A partitioning of the graph is optimized until a local maximum of the modularity is found [12]. A noted weakness of this approach is its inability to detect communities of varying size [13] [14]. This problem, termed the resolution limit, is caused by small communities being merged and large ones split. This is of consequence to scRNA-seq analysis, as cell types vary in abundance by orders of magnitude [15].

Multi-resolution measures of modular structure, parameterized by the resolution parameter *γ*, were introduced to control the sizes of the partitions identified [7]. While these quality functions give control of the sizes of the partitions to the analyst, they aren’t able to find large and small communities in the same solution [16]. This raises the question of how the parameter is selected, and what can be learned from the structure at different scales. One approach is through stability of the solution – where a good parameterization produces consistent partitioning solutions for small changes in the parameter [17] [18]. By scanning a range of parameters, these methods can find partitionings of the network at multiple scales, but can only identify global solutions. This precludes the identification of groups which are only stable when other areas of the graph are not, and can only be found by individual assessment. We would expect these groups when dealing with complex community structure, like those found in large single cell or other biological data sets. For example [19] uses a scan of the resolution parameter to describe modules within protein-protein interaction networks at different scales. This method looks at how modules of protein change with resolution, but do not use stability to identify discrete modules. Within the single cell literature, the `clustree` tool ([5]) provides visualizations of how cluster identity changes as parameters vary, which can allow subjective identification of stable groups. By viewing how clustering solutions change along a range of parameters, stably identified ones can be identified as regions of the tree with few edges.

While these methods demonstrate how varying a single parameter of a clustering method can be used to examine the structure of a data set, scRNA-seq analysis pipelines frequently have many parameters. Approaches to find robust clusters under uncertainty of multiple parameters generally falls under the area of consensus clustering. Consensus based methods have been used for scRNA-seq by SC3 [20] and RSEC [21]. In each of these methods, a data set is clustered many times and single partitioning which best fits with each of the solutions is generated. The approaches don’t model the effects of individual parameters on the resulting clustering, and so don’t allow for identification of multilevel solutions based on parameter choice. Here, we propose to formalize and extend these operations by automatically detecting the clusters which are consistently found within contiguous regions of parameter space.

## 3 Methods

### 3.1 Overview

Our underlying assumption is that a true cluster of samples should be robust to parameterization of the clustering algorithm [22]. Previous techniques, like tight clustering [23], have used this intuition to iteratively find individual stable clusters, but can only find a single level of structure in the data. Traag et. al.[18] finds stable global solutions when a single parameters is varied. However, typical processing of scRNA-seq leading up to unsupervised clustering involves many parameters [24]. In addition, we expect multiple clusters at different resolutions would reflect the biology better than a single global solution. `constclust` implements a novel approach to identifying and prioritizing clusters to address these issues.

First, the parameter space of the clustering method is explored in a grid-search like strategy, Fig. 2 (a). The parameters being varied must have an ordered relationship or be equivalent to random restarts. This is necessary so we can have locality in parameter space: we need to be able to say which settings are close to each other to have any expectation about their effect.

**Figure 2:**
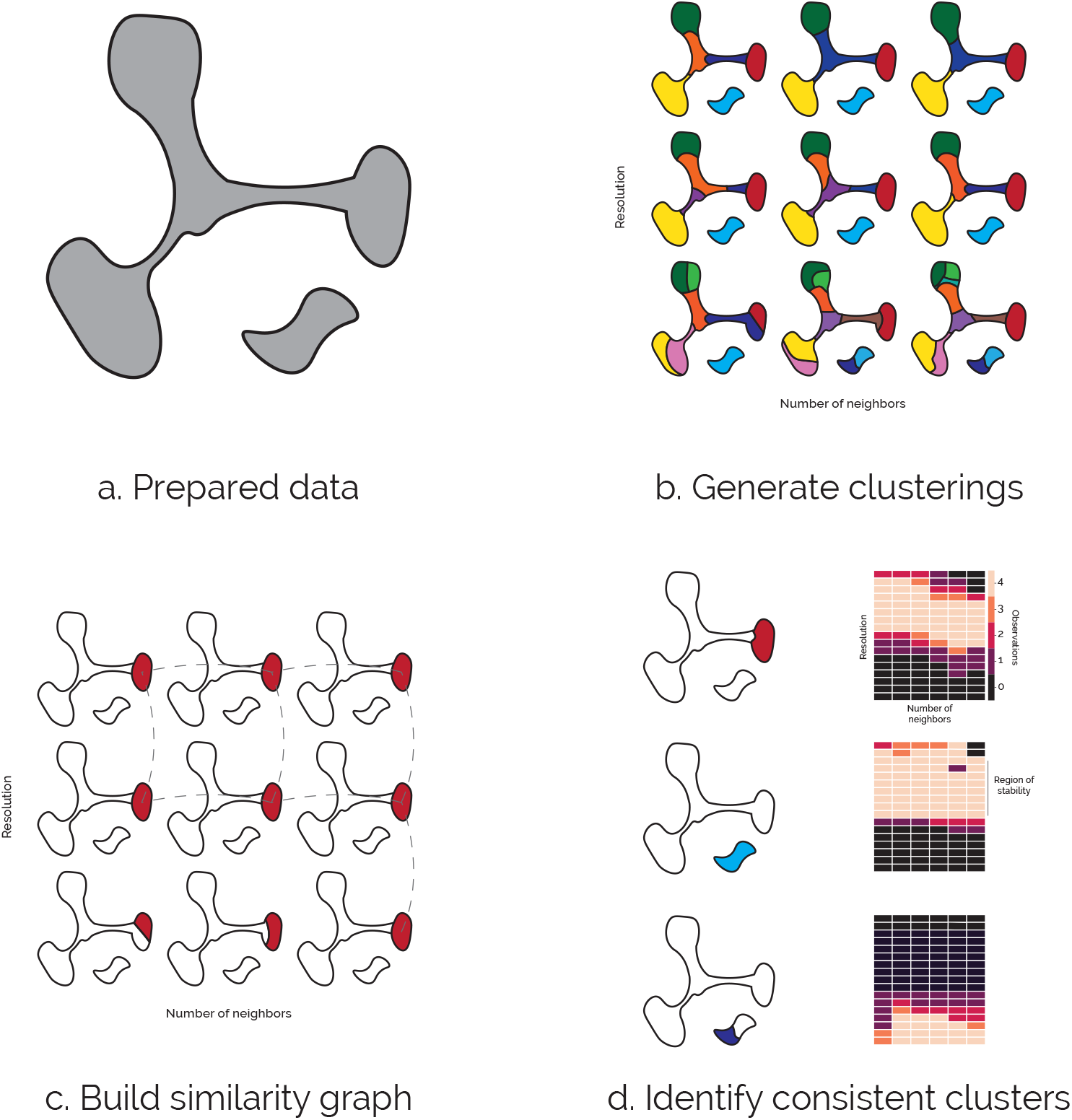
An overview of the constclust method. (A) Preprocessed data (the feature space which clustering will performed on) is partitioned across a range of parameters, (B) using a gridsearch like sampling strategy. (C) A similarity graph of the identified clusters is then built, where edges are the Jaccard index of clusters from neighboring partitionings. Low weight edges are pruned from the graph and the remaining connected components are collected. These components correspond to a consistently found cluster. (D) As each node within the component is a cluster found from a run of the clustering method, the parameter region it was identified in can be derived.

Clusters which are found over a wide range of parameters are likely of high quality. We return the set of unique (within a similarity cutoff) clusters identified ranked by the size of the parameter space they were found in. This creates a prioritized list of clusters for the cells, in contrast with the more common flat partitioning result. We believe this more closely with our concept of cell-types or communities, since we don’t assume there is a single set of disjoint labels which cover all samples. Rather, the samples are classified with a set of labels which can be hierarchical or overlapping.

Additionally there is no promise every (or any) individual sample is assigned to a stable solution, as can be expected in data with continuous states.

### 3.2 Implementation

#### 3.2.1 Generating clusters

The input of `constclust` comprises of the data set (expression matrix) and a clustering method. It then performs clustering on the data set across a range of hyperparameters in a grid-search like strategy. Each clustering solution specifies an assignment of each sample to a cluster. While any clustering method and set of parameters could be used, constclust comes with a convenience method for efficiently generating modularity based clustering based on the typical `scanpy` workflow [8]. Modularity based clustering was chosen due to it’s performance within the field [11], that there is pre-existing literature for connected approaches, and since it is quick to run.

In the default implementation, the weighted graph of nearest neighbors is computed using UMAP [25] on the top 50 components of a PCA of the data set. This graph is then partitioned using the `Leiden` optimization algorithm [26] using Reichardt and Bornholdt’s Potts model [27] as the quality function. The parameters varied are the number of neighbors used to construct the graph and the resolution metric for the quality function. We consider these to be ordered parameters, since we can say 45 neighbors is closer to 30 neighbors than 60 neighbors is. Because of this we have can have steps in these sets of parameters. For example, given the resolutions `[0.5, 0.6, 0.7]`, the resolution `0.6` is one step from `0.5` and `0.7`, while `0.5` is two steps from `0.7`. Partitions separated by one step in the ordered parameter grid are called ‘neighbors’ in parameter space.

For robustness of results, the data set is clustered multiple times with each of these ordered parameters using different random seeds to initialize neighbor finding. Random seeds are considered an unordered parameter, as we don’t expect solutions from setting a seed to 5 to be any more similar to those generated with seed 3 than those generated with seed 1. This reflects a noise-based approach to resampling the partitionings, as opposed to a bootstrap approach (as used by [21] [28]).

#### 3.2.2 Comparing clusters

Cluster-wise comparisons are made between neighboring partitionings. An undirected weighted graph is built from these comparisons, *G*(*n, e*) where nodes *n* are individual clusters and edges *e* is weighted by Jaccard similarity of the clusters. Two clusters which contain identical groups of samples will be connected by an edge of weight one, while clusters with no overlap will have an edge of weight zero (equivalent to no edge) [29].

#### 3.2.3 Resolving stable clusters (ranked list)

The edge weight distribution of the resulting graph is bimodal (supplementary), with a large number of high weight edges and near zero edges. This suggests a consistent core set of clusters. To find those, allowing for some fluctuation in identity, we remove low weight edges using a user specified cutoff and identify the remaining connected components of the graph. From the components of this graph we can find the range of parameterizations for which any one consistently found cluster could be identified, and how the samples within it change.

Each of the components represents a set of cells which were consistently clustered together over a contiguous region of parameter space. While selecting components to use in downstream analysis is now up to the user, `constclust` provides tools for identifying components to use in downstream analysis. For visualization, a view of where the samples sit in a reduced dimension plot, and the region of parameter space can be visualized (Fig. 2 (d)). As cells can exist in multiple components, hierarchies of components which share samples can be inferred. This structure is presented to the user as a tree where each component is a node, and any larger component it’s cells also are found in is that node parent. Users can visualize the component and see some summary statistics via hover over on these plots (Fig. 5).

### 3.3 Data processing

#### 3.3.1 Simulated data

Simulated data sets were generated using `splatter` [30] with default parameters. For the clusters of equal sizes, the data set consisted of 5000 samples with four true clusters of equal size. When varying cluster sizes 10,000 samples were generated, with four true clusters with group probabilities of: .9, .09, .009 and .001.

#### 3.3.2 PBMC dataset 10x

Aligned PBMC expression data was retrieved from 10x genomics https://support.10xgenomics.com/single-cell-gene-expression/datasets/3.0.0/pbmc_10k_v3, initially consisting of 11,679 cells annotated to 33,538 genes. This data set has been preprocessed and filtered for doublets (9,936 cells left after preprocessing) before having the constclust workflow run with it. For this data set 1800 separate clusterings were generated and resolved into components which showed up in at least a hundred solutions.

#### 3.3.3 Resampling

To assess the robustness of components identified with constclust the data set was subsampled and the results compared with the full run.

Sample frequency vectors were calculated by counting (for all cells) how many times each showed up in a component across all of it’s clusterings. These were normalized by total number of clusterings in a component – so a cell which showed up in every clustering would have value 1, and a cell which showed up in no clusterings would have value 0. To find the nearest solution in one of the resampled data sets, each cell weighting vector was subset to have only the cells present in each data set, and pairwise cosine distances between the components of each data set calculated.

### 3.4 Metrics

#### 3.4.1 Geary’s C

Geary’s C is a method for determining local auto-correlation in networks [31]. This metric measures, for some value on each node of a graph: are those values correlated with that node’s neighbors’ value.

This computed as:

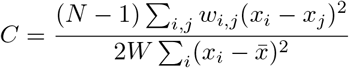

Where *N* is the total number of nodes, *w_i,j_* is the weight of the edge between nodes *i* and *j*, *x_i_* is the value at node *i*, and 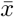 is the mean value of *x*, and *W* = Σ*w_i,j_*.

This method is typically used in spatial contexts, to determine if a measure is correlated with it’s location [32]. It has also been used in scRNA-seq analysis to determine whether a feature was correlated with transcriptome similarity in the `VISION` tool [33].

### 3.5 Code and data availability

`constclust` is available on pip (`pip install constclust`). Source code is hosted at https://github.com/ivirshup/constclust. while documentation and tutorials are available at https://constclust.readthedocs.io. Supplementary information and code to reproduce the analyses used in this manuscript are available as Jupyter notebooks at https://github.com/wellslab/constclust_supp along with instructions for retrieving the data.

## 4 Results

### 4.1 Planted clusters are clearly identified

To demonstrate some of the basic concepts behind `constclust`, we’ve applied it to simulated data. First, it’s run on a data set with four groups with even sizes (Fig. 3). Unsurprisingly, these are easily identifiable by constclust, as they can be easily identified by the underlying clustering algorithm. For the majority of the parameterizations tested, the correct underlying representation can be found exactly. While `constclust` finds labels which match ground truth, so does running the underlying clustering method with default parameters. This demonstrates the true solution is robustly found while false solutions are clearly separated.

**Figure 3:**
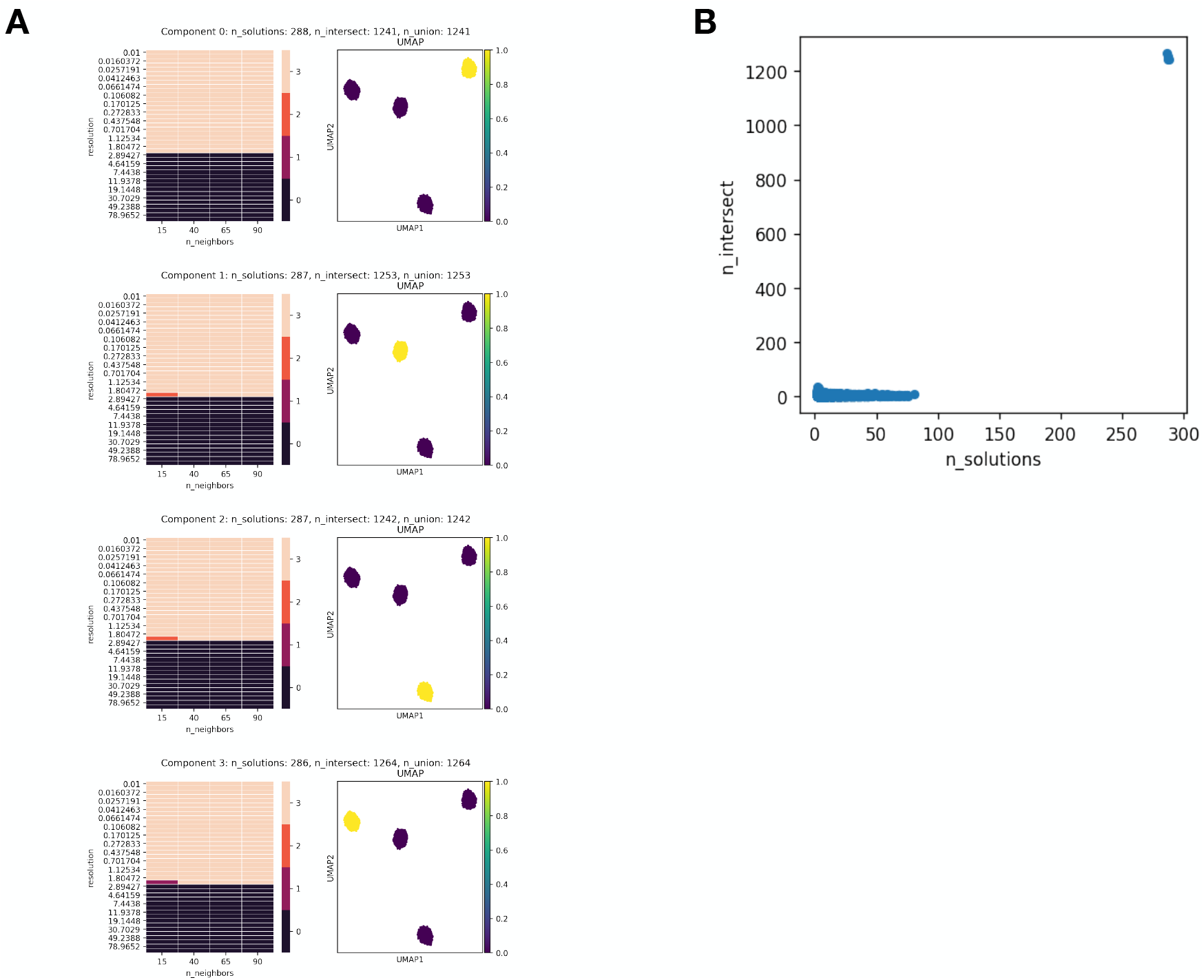
Results of constclust on a simulated data set of 5000 cells with four equally sized clusters. This data set was generated with Splatter, and was then clustered 480 times varying number of neighbors, resolution, and with four random initializations per parameterization. The correct assignments of cells to cluster show up in the top four components (a) found by constclust. (b) These components showed up in a minimum of 284 solutions. These components clearly stand out from the other solutions found by both their size and frequency of occurrence, e.g. the four selected components are in the top right. The next most common clustering solution showed up in 84 solutions and contained only 8 cells. Notably, a solution containing all of the correct groupings would show up in both the standard parameterization, as well as a number of other solutions. All that is gained here by using constclust is a greater confidence in the solution from the clustering algorithm.

### 4.2 constclust can detect planted clusters of vastly different scales

As cell types occur at widely varying frequencies, constclust was next tested against a data set whose underlying groups varied across four orders of magnitude (Fig. 4). This simulation is a more realistic depiction of cell type frequency. Notably, there was no one clustering solution which correctly groups both the smallest and largest planted cell types together. Instability of solutions when no stable community can be found follows from expectations from the community detection literature.

**Figure 4:**
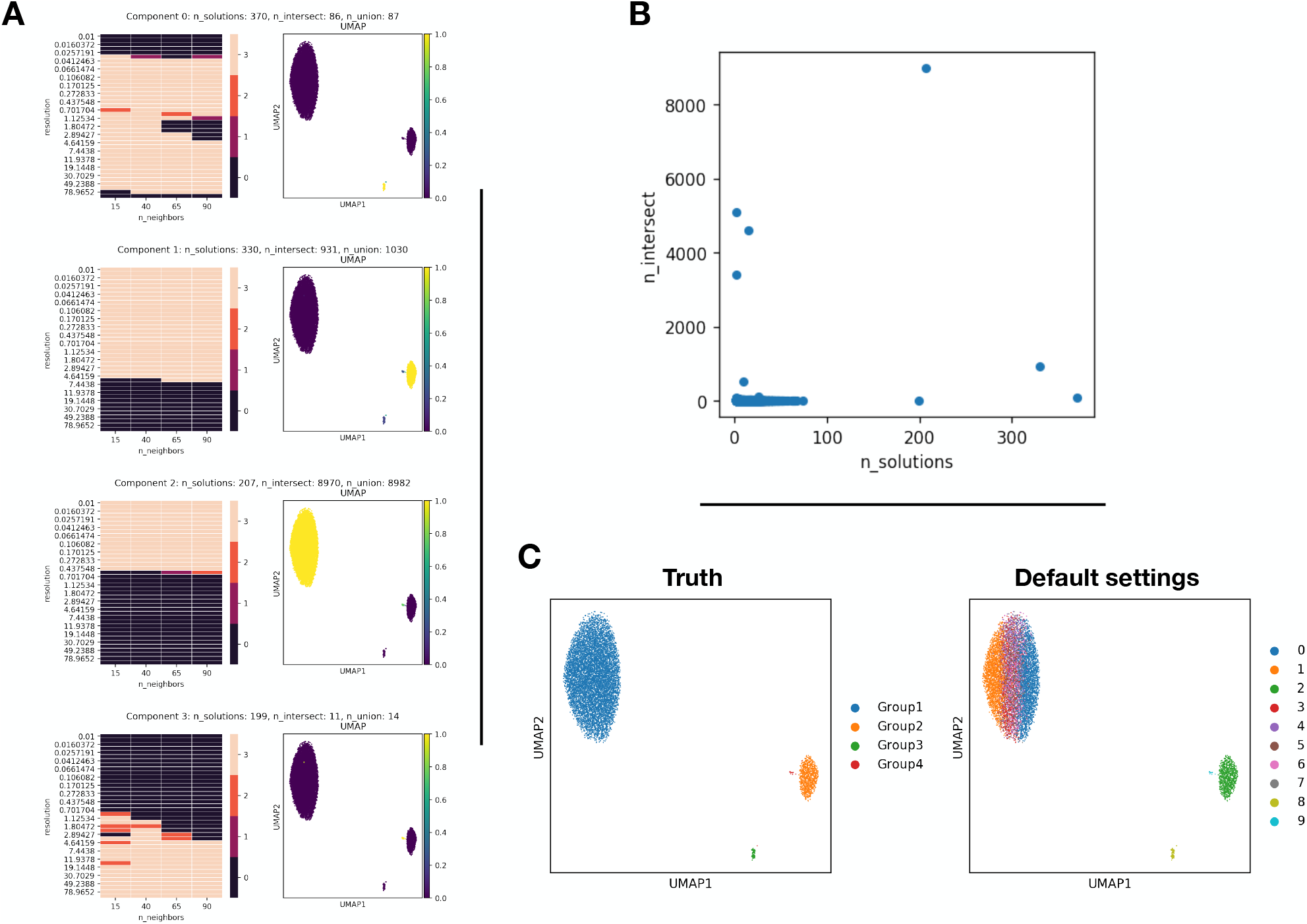
Results of constclust on a simulated dataset with clusters of varying sizes. The dataset here consisted of 10,000 simulated cells, and has four planted clusters which vary in size across orders of magnitude. (a) shows the top four components, and the range of corresponding parameters. These components match the planted clusters with an ARI (adjusted rand index) of 0.999. (b) Similarly to the evenly sized clusters, components corresponding to true clusters were distinctively more common. (c) Notably, there is no single solution which contains each of the correct labellings. As expected from the community detection literature, the largest cluster randomly splits at resolutions where the smallest cluster can be detected.

### 4.3 constclust reveals multilevel structure in single cell data sets

Instead of having a single complete labelling for a data set, `constclust` generates a collection of labels (clusters), each of which apply to a subset of the samples. This re-frames the problem from partitioning a data-set with a single cover, to finding a set of computationally robust labels which represent real biological signal. One key features of this structure is that individual samples can be assigned more than one label. We demonstrate this on a data set of ten thousand PBMCs from 10x genomics. We use this structure to assign cell labels at a high level of lineage (Fig. 5 (a)) as well more specific functional categories (Fig. 5 (b)). Using the set of stable components found in this data set, we were able to create an interpretable labelling of the samples.

**Figure 5:**
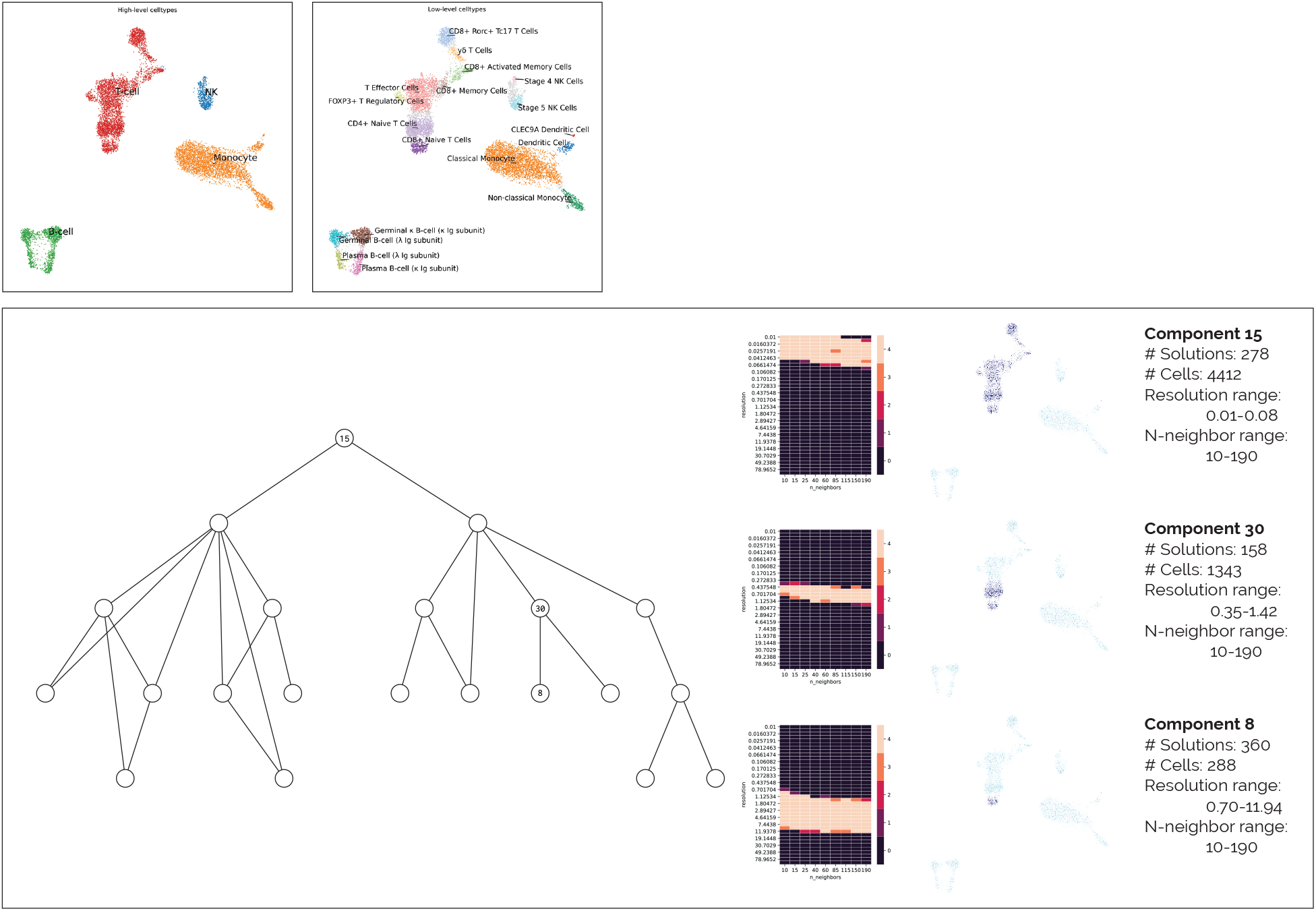
Multilevel structure identified by constclust. A high and low level clustering are shown in a and b respectively. The high level clusters define high level populations of monocytes, natural killer cells, B-cells and T-cells. More fine grained populations (*e.g.* cytotoxic vs non t-cells) can be found by looking at the smaller components. c shows a hierarchy of clusters found in the leukocyte population. Each cluster here was identified in at least 10% of solutions 1440 solutions total. In the hierarchy, a component is considered the child of another component if it is smaller and contains a subset of the cells. On the right we show which groupings of cells were identified by plotting and the range of parameters they were identified for. This visualization is available as an interactive plot in the constclust package. Though components were not manually curated, each here shows at minimum 273 differentially expressed genes (*P* < 0.5 Mann Whitney U test with BH adjustment) when compared against the rest of the data set.

To create the kind of flat labeling seen in Fig. 5, 23 components out of a broader set of stable solutions were selected. This is a reduced view of the signal we were able to identify, and required manual curation to collate. A hierarchical view can be built from stable components based on the overlap of their samples. In the PBMC data set we can find three levels of identity for CD8+ Memory T-cells. At a high level, we can see the broader T-cell population (component 15), followed by naive T-cells (component 30) (CCR7+, CCL5−), finally resulting in naive CD8+ T-cells (component 8) (CD8+, CCR7+, CCL5−) (Fig. 5 (c)). This provides an intuitive interface for users to explore the structure of their data-set at multiple scales of resolution by “drilling down” from broad to specific communities.

There are a few key differences between the approximate hierarchy shown here and those created by more strictly hierarchical or agglomerative methods (like those used by `clusterExperiment` [21] or `dendrosplit` [34]). As each component has can be identified independently of the others, there is no restriction that components must be strict subsets or supersets of other components. This also means that there does not have to be a set of components which contains each sample exactly once, a restriction of common clustering methods like K-means or modularity minimization. When selecting the sets of components for a flat labelling there will be cells left unlabelled if they didn’t clearly fit into a discrete group.

### 4.4 constclust robustly identifies discrete groups of samples

To assess the quality of the components identified by constclust, we used three methods. The first is the intuitive metric of ranking components of samples by the number of clustering solutions they are found in. Highly ranked components were found in more clustering solutions. Here we show that these highly ranked components are picking up true signal in the data through a re-sampling experiment and measuring their autocorrelation on a graph representation of the data set.

If we are able to identify true sub-communities of cells in a data set, we would not expect those communities to change if only a subset of the data is measured. To show this, `constclust` was run 280 times on subsets of the previously used PBMC data set (Fig. 6). The subsampled results were searched for their closest matching component from the original set. This forms a clearly bi-modal distribution of distances where the more highly ranked components are consistently matched. High ranking components were consistently found between runs. However, due to decrease in data set size, top ranked components in the full data set with small cluster sizes were often dropped in the subsampled data sets. While some lowly ranked components (126, 137, 142, 143, 151, 182) from the original assignment appear to have good matches in the permuted data-sets. In most comparisons, there was a better match at a lower rank (supp.).

**Figure 6:**
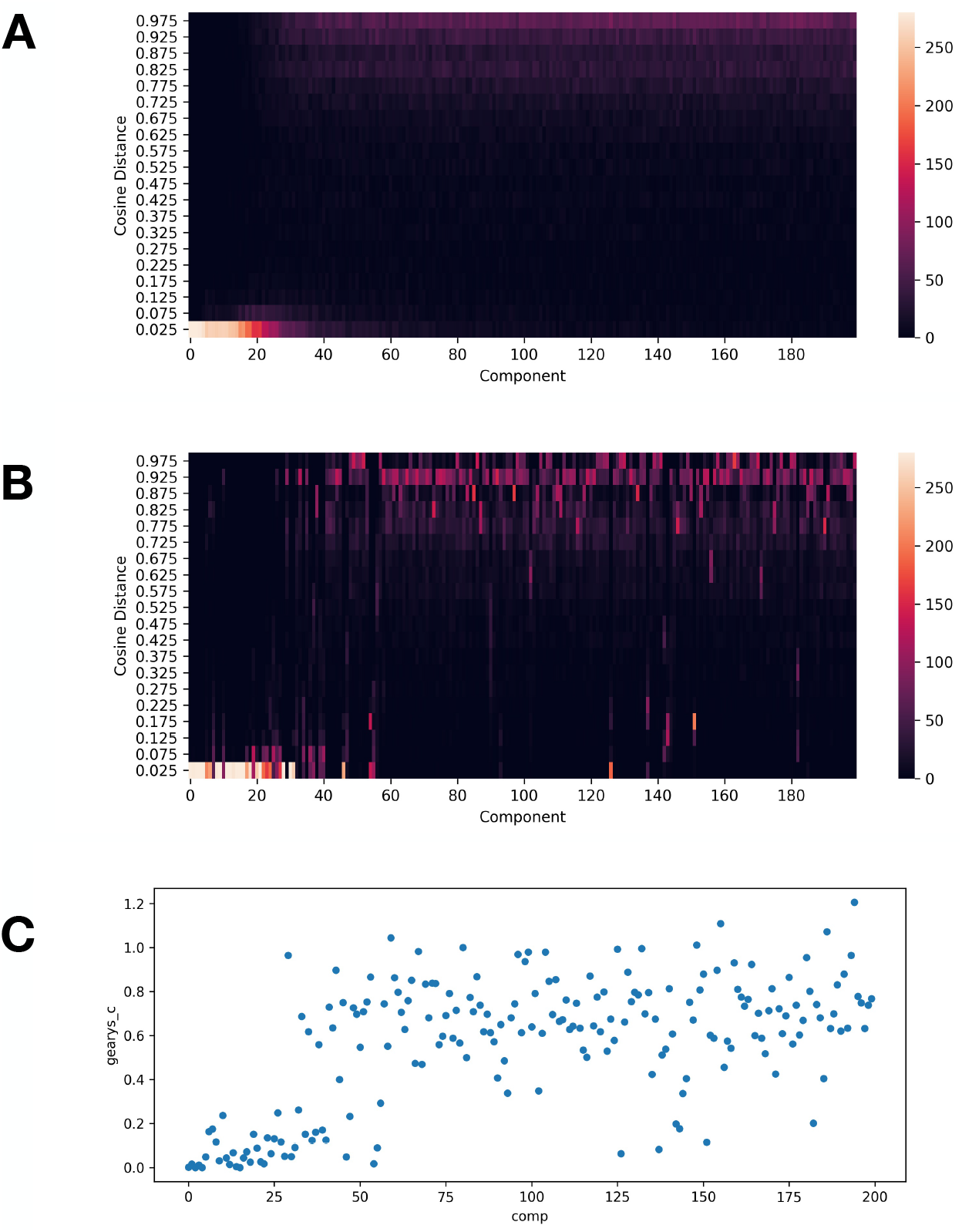
Highly ranked components are consistently found in resampling experiments. To assess the stability of the components and their rank, a re-sampling experiment was performed. The component _nding pipeline was run again on 280 randomly selected subsets, each 1/2 the original data sets size, of the PBMC data set. The cosine distance of cell frequencies for each pair of components between the top 200 ranked components in each the resampled and original set. (A) shows the distribution of the minimum distance to each components closest match (y-axis) by the components rank in the subsampled clustering (x-axis). Components are also accurately identifying discrete communities in a KNN representation of a data set. (C) For the PBMC data, Geary’s C (yaxis was calculated for the cell frequencies of each component and plotted against that components rank (x-axis).

To assess whether identified components are real signal in the data, we used the Geary’s C metric to measure how well a component explained variance in a nearest neighbor representation for the data set. Previously, this metric has been used by `VISION` [33] to find pathways which are spatially discriminating in a latent space. Here, we expect samples from the same community to be near to each other. That is, we expect the sample labels for a cell in the KNN graph to be correlated with their neighbor’s labels, and given a low value of Geary’s C. This is very useful for discriminating between signal and noise in our data sets, as noise should be randomly distributed. And indeed this is what we see Fig. 6 (c) there is a bimodal distribution of these values.

### 4.5 Computational Efficiency

Compared to other consensus clustering frameworks for scRNA-seq, `constclust` is highly performant (Table 1). This is due to the use of a fast clustering algorithm and an embarrassingly parallel formulation which allows near-linear scaling with available cores. Work to improve this performance is ongoing, with an aim at reducing time spent distributing the data. Another source of increased performance is that `constclust` does not compute a dense coincidence matrix like SC3 and RSEC, resulting in memory use of *O*(*cells* × *partitionings*) as opposed to *O*(*cells*^2^). The performance advantages these approaches provide is clear when compared with other methods. For a data set of 12 thousand cells, `constclust` could generate and reconcile 4,500 partitionings in less than half the time SC3 (running on a subset of the data, and classifying the rest with an SVM) takes.

**Table 1:** Run times and number of clusters generated for consensus clustering methods designed for scRNA-seq data. All results are based on a 12 thousand cell data set using on 10X’s v2 chemistry. SC3 [20] was run with its default parameters, meaning it ran its standard algorithm on five thousand cells, then classified the rest using an SVM. RSEC [21] was run with default parameters, and had it’s process killed after running for 36 hours.

|  | constclust | SC3 (SVM mode) | RSEC |
| --- | --- | --- | --- |
| Time | 2.5 hr | 6.8 hrs | ? (>36 hrs) |
| # of partitionings | 4500 | 150 | ? |

## 5 Discussion

The ability to identify cell types and states from single cell high-throughput assays depends on identifying meaningful clusters of cells with unsupervised methods. That is, the classification of cells by scRNAseq markers, or identification of new cell phenotypes relies first on accurate groupings of those cells. Increasingly we see that complex biology can’t be captured in a single ‘flat’ clustering solution, but requires analysts to navigate hierarchies captured within the data. ‘constclust’ provides a new way to approach the problem of finding the right clustering solution.

`constclust` was developed to address the issues of parameter choice and class imbalance when clustering single cell data. By aggregating clustering solutions using locality in parameter space we find clusters that are consistently identified. In data sets with planted communities, `constclust` specifically found ground truth labels with high accuracy by evaluating their stability over a range of clustering parameters. Additionally, constclust could find planted communities even when they vary by orders of magnitude and could not be captured by one set of hyperparameters. When applied to real-world data, known cell hierarchies and types were readily identified. Finally, approaching the method by looking for local instead of global stability allows for significant reductions in compute time and memory usage over other consensus approaches.

The observation that ‘flat’ clustering solutions don’t typically capture biological complexity is a known issue in the field [6] [4]. Most popular clustering methods only find flat clusterings of the data set. This does not fit with the goal of finding cell types and cell states present, possibly across biological hierarchies.

Parameter choice is increasingly recognized for the huge effect on the results of unsupervised clustering for single cell data, however selecting these values is still an open question [35] [5] [36]. Recent approaches to find robust clusterings like those from [28] [37] look across many clusterings generated with different hyperparameters, but only provide tools for evaluating the paritionings as a whole. As we show with simulated data, some planted communities cannot be detected with a single choice of hyper parameters. By looking at the literature from the fields of community detection, we take a different view based on the idea that clustering algorithms may not be able to find a good solution for all samples in the data set at once. Using this view, we look only for individual stable clusters – which fits much better with the multilevel structure of cell populations.

The idea of finding a better clustering solution by combining the results of many individual solutions from different parameterizations is not new. A typical consensus clustering approach (like those used by [20] or [21]) of looking at the space of generated cluster solution and determining a likely single true clustering. These consensus clustering methods are asking if a common or average single clustering solution can be found from a set of input solutions [38]. While `constclust` similarly uses information from multiple clustering solutions to generate a consensus categorization, it asks a different question. Instead we look at the space of generated clusters and try to filter out the unstable ones by asking “am I as similar to my neighbors as they are to theirs”.

`constclust` stems from the same intuition that Evidence Accumulation Clustering (EAC) is based on. In EAC, each clustering solution is considered an independent piece of evidence about the natural partitions present in the data. If two samples are put in the same cluster, this is evidence those samples truly are part of the same group [39]. In `constclust`, we extend this. First we recognize that if the set of clusterings to be ensembled are generated by varying parameters, the solutions are actually dependent on those parameters. Additionally we do not look for evidence that two sample sit in the same cluster, instead we are looking for evidence that the entire cluster is a true grouping of the data.

A similarity can be drawn here to density based clustering or outlier detection. These methods take local context into account when comparing objects [32] – though the definition of context may vary. For example, when we make a KNN representation of our data the context is the K most similar samples in our data set. In `constclust` we use two kinds of context for each cluster, (1) the surrounding parameter space and (2) clusters with a shared set of cells. From this definition of locality, we can ask which clusters are like each other within a certain context. A random grouping of samples should be an outlier, since it isn’t like any other sample within this context. In this way, the problem is outlier detection, similar to methods like HDBSCAN [40]. Once we’ve identified the stable solutions with the clustering space, we have identified our components.

### 5.1 Considerations for evaluation

One of the key features of `constclust` is that it returns a set of labels. This contrasts with methods which return either flat or hierarchical solutions. The shortcomings with flat solutions have been extensively discussed and recognized within both clustering and single cell analysis [4] literature. However, returning a different kind of structure makes comparison difficult.

While metrics like Adjusted Rand Index (ARI) or cluster silhouette are commonly used when assessing performance of clustering method for scRNA-seq, they can’t be used here. Both ARI and silhouette require a single complete labelling of the data. In addition, ARI requires a ground truth labelling for the data to compare against. This is problematic here `constclust` does not generate flat labellings and since multilevel labellings of data sets are rare. We suggest the intuitive quality metric of simply ranking components of samples by the number of clustering solutions they are found in. This approach is validated by results showing highly ranked components are picking up true signal in the data through a re-sampling experiment and measuring their autocorrelation on a graph representation of the data set.

Unlike other multilevel methods like `TooManyCells` [41] and `HDBSCAN` [40], `constclust` has no restriction of strict hierarchical structure being found in the data. In hierarchical models, the samples which can be found in a partition are dependent on other partitions in the data set. They must be complete subsets or super sets. This is an important feature, since we don’t expect biology to behave like this. For example, when attempting to perform a cross species data integration, what is the hierarchy to be used? When macrophages can be derived from different lineages, which hierarchies should they sit in [3]? If a hierarchical structure is assumed, these kinds of relationships are difficult to model.

Assessment of non-flat or non-hierarchical clustering methods is an open topic for scRNA-seq analysis. Since the problem is foundational to the goal of cell-type discover, it’s critical this be addressed. Recent work to this end include new metrics for evaluating hierarchical clustering solutions [42]. The development of gold standard data sets with multilevel annotations, and further methods for evaluating multilevel solutions is an important direction for the field.

### 5.2 Limitations

There remain a number of limitations with `constclust`, and challenges for the field more generally.

While multiple levels of structure can be found, there’s no particular reason these must correlate with a biologically distinct group of cells. This can only be confirmed through downstream analysis using outside knowledge. Of course applies to any unsupervised clustering method using scRNA-seq data, since the mapping from transcriptome to cell type is by no means resolved. However, this does not fit the structure given by other analysis tools.

All analyses in this manuscript rely on cutoffs, for determining which components are worth investigating further. While future work could expand into better prioritization of components, this is alleviated by some tooling. For example, for the classical monocytes from the PBMC dataset the next stable solutions (occurring in at least 100 clustering solutions) which can be identified from that group of cells have a maximum of 7 cells (compared to the 2500 monocytes) and show no differentially expressed genes below a *p* ≤ 0.05 cutoff in a pairwise comparison using a Wilcoxon test.

Nor do we completely relieve the analyst of the burden of parameter selection. While schemes for automatic exploration of parameters could be defined, this would be dependent on the clustering method and parameter. Additionally if we know what a good parameter space was for our clustering, then this method probably wouldn’t be needed. What ‘constclust’ does provide is a straight forward way to scan across parameters, and identify stable clusters at different data scales, to allow a user to find subgroupings of cells that might otherwise be hidden in a single global ‘best fit’ solution.

## 6 Funding

KALC was supported by the National Health and Medical Research Council (NHMRC) Career Development fellowship (GNT1159458). JC is funded by NHMRC (GNT1181327) and (APP1186371) to CAW https://www.nhmrc.gov.au. IV is funded by the Centre for Stem Cell Systems https://biomedicalsciences.unimelb.edu.au/departments/anatomy-and-neuroscience/engage/cscs The funders had no role in study design, data collection and analysis, decision to publish,or preparation of the manuscript.

## Footnotes

1 <https://github.com/ivirshup/constclust>

## Notes

https://github.com/ivirshup/constclust

https://constclust.readthedocs.io

https://github.com/wellslab/constclust_supp

### Competing Interest Statement

The authors have declared no competing interest.

